# Operations Research Methods for Estimating the Population Size of Neuron Types

**DOI:** 10.1101/633313

**Authors:** Sarojini M. Attili, Sean Mackesey, Giorgio A. Ascoli

## Abstract

Understanding brain computation requires assembling a complete catalog of its architectural components. Although the brain is organized into several anatomical and functional regions, it is ultimately the neurons in every region that are responsible for cognition and behavior. Thus, classifying neuron types throughout the brain and quantifying the population sizes of distinct classes in different regions is a key subject of research in the neuroscience community. Although the total number of neurons in the brain has been estimated for multiple species, the definition and population size of each neuron type are still open questions even in common model organisms: the so called cell census problem. We propose a methodology that uses operations research principles to estimate the number of neurons in each type based on available information on their distinguishing properties. Thus, assuming a set of neuron type definitions, we provide a solution to the issue of assessing their relative proportions. Specifically, we present a three-step approach that includes literature search, equation generation, and numerical optimization. Solving numerically the set of equations generated by literature mining yields best estimates or most likely ranges for the number of neurons in each type. While this strategy can be applied to any neural system, we illustrate its usage on the rodent hippocampus.

## 1 Introduction

A quantitative description of the brain’s machinery is essential to understand the mechanisms of nervous system functions. The brain encompasses an extraordinary quantity and diversity of cells. The human brain contains around 100 billion neurons (Herculano-Houzel 2009) and the rodent brain contains around 100 million neurons (Herculano-Houzel et al. 2011). Neurons can be grouped into many distinct types based on their structural, physiological and molecular features (Bota and Swanson 2007). The composition of balanced proportions of neuron types into elaborate networks enables the brains many specific computations. Estimated counts of neuronal types, i.e. a “neuronal census”, would enable more accurate and complete models of brain circuits. Towards this goal, the National Institutes of Health launched the BRAIN Initiative Cell Census Network (BICCN), a consortium of research projects tasked with generating a comprehensive molecular and anatomical cellular parts list within a three-dimensional reference mouse whole-brain atlas (Ecker et al. 2017).

Counting the neurons of each type in a region requires establishing the identity of millions of individual neurons. Rapid progress in genetic phenotyping is on the verge of enabling a comprehensive cell-level classification of neurons throughout the mouse cortex (Tasic et al. 2018). However, linking this growing molecular data to anatomical connectivity requires the analysis of the neuronal input and output elements, namely dendritic and axonal arbors. Full morphological characterization of axons and dendrites involves physical or optical tissue sectioning to follow the complex branching structures in the dense three-dimensional space. This is a labor-intensive and error-prone procedure for a human to perform manually, underscoring the need for increasingly automated machine-learning approaches (Peng et al. 2015; Januszewski et al. 2018). Experimentally, the problem is exacerbated by the large disproportion between the total length of an individual axon (hundreds of millimeters) and its branch thickness (tens of nanometers), resulting in a very small ratio (~ 10^−7^) between the physical space a single neuronal projection overall invades and the volume it actually occupies. This major obstacle will likely keep the acquisition of comprehensive structural data at single-neuron resolution below full-brain scale for many years. Therefore, indirect estimation of neuron type population counts is an important and useful endeavor.

The neuroscience literature contains a great deal of data relevant to the census problem. These include stereological sampling of neuronal densities in specific anatomical areas, morphological characterizations of collections of neurons from the same brain region, slice imaging of neurons stained for a particular molecular marker, and more. Each of these data types expresses facts about absolute or relative neuronal population sizes. Harnessing such diverse sources of information towards a neuronal census poses two primary challenges: (i) formatting all relevant observations in terms of a common neuronal classification scheme; and (ii) inferring population sizes from the properly formatted evidence. Solving the first challenge will ultimately require a broad consensus in the neuroscience community on how to define neuron types objectively and reproducibly (Armañanzas and Ascoli 2015). For the purpose of illustration, in this study we tentatively adopt a recent circuit-based classification proposal (Ascoli and Wheeler 2016) for which relatively abundant data are available for parts of the rodent brain such as the hippocampus.

Solving the second challenge entails a workflow to integrate possibly contrasting measurements while interpolating through missing data points. Operations research offers many techniques for leveraging inconsistent and/or incomplete information to achieve an optimal estimate for a set of target parameters. These techniques fall under the broad umbrella of *mathematical optimization*. Here we describe the use of mathematical optimization to obtain an estimated neuronal census. The neuronal population counts to be estimated are represented as free parameters. Data relating neuron types to their properties (e.g. from literature search or experiment) are formatted as equations in terms of these parameters. These equations are composed into an objective function, which can be optimized by a variety of algorithms. Below we present a brief abstract overview of relevant mathematical optimization methods and a concrete application to the hippocampal subregion of the dentate gyrus.

## 2 Methods

In order to describe the operation research aspects of our approach, it is useful first to explain how it is possible to derive a system of equations representing useful constraints regarding a neuronal census. In the most general sense, every neuron type is associated with a distinct collection of properties (e.g. morphological, physiological or molecular) through a many-to-many relationship. In other words, no single property uniquely identifies a neuron type, and any property is typically associated with multiple neuron types. However, the full set of properties of a given neuron type is indeed different from that of all other neuron types. Useful constraints consist of measurements, observations or reports on neuronal properties that can link combinations of neuron types to specific numerical values.

Consider for instance a brain region with only two neuron types, A and B, and corresponding counts *n*_*A*_ and *n*_*B*_. If a stereology experiment determines the total number of neurons in that region to be 1000, this provides a useful constraint (and corresponding equation) by setting the sum of the two target counts to the measured value (*n*_*A*_+*n*_*B*_ = 1000). Now suppose that only neuron type A expresses a particular protein and an article reports that, out of 20 cells tested in that region, 15 were found to be positive for that protein while 5 were negative. This provides another useful constraint (and a second equation) by indicating a ratio (3:1) between the two target values 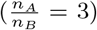. In this simple case the number of independent equations equals the number of unknowns yielding a well-constrained system with a single exact solution (*n*_*A*_ = 750, *n*_*B*_ = 250).

In a more general sense, a system of equations is overdetermined if there are more constraints (independent equations) than parameters and underdetermined if there are fewer constraints than parameters. In the census problem, overdetermined system may arise from multiple experiments measuring the same variable (e.g. the total number of neurons in a region) and yielding different results. As overdetermined systems are inconsistent, they do not have exact solutions. In this case numerical optimization may find an optimal estimate that minimizes the discrepancy from all available constraints. Underdetermined systems arise when insufficient constraints are available for one or more of the target unknowns. If none of the constraints are mutually inconsistent, an underdetermined system typically has an infinite number of solutions. In this case numerical optimization may find the range of values defining the possible solutions.

To illustrate the optimization problem we use the following example. Suppose there are five unknown neuronal population counts represented by parameters *X*_1_, *X*_2_, … *X*_5_, and the known facts about these populations define the following system of equations:

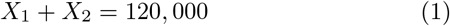

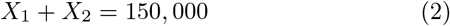

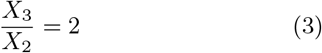

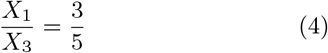

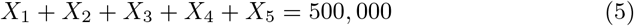

Although the number of equations equals the number of target unknowns, parameters *X*_1_ and *X*_2_ are over-constrained because equations (1) and (2) are inconsistent, whereas parameters *X*_4_ and *X*_5_ are under-constrained and therefore will admit a range of solutions.

Numerical optimization starts by constructing from the system of equations an objective function to be minimized (or in some cases maximized). If the system contains non-linear equations, the objective function is also nonlinear, whereas either linear or non-linear objective functions can be constructed from linear equation systems. A common objective function consists of the sum of the weighted squared errors (differences between the two sides) associated with each equation. Errors can be weighed in accordance with the relative reliability of the data from which the equation is derived. Suppose, for instance, that equations 1, 2, 4, and 5 in the above example were considered to be 4 times more reliable than equation 3. Then the objective function to be minimized would be:

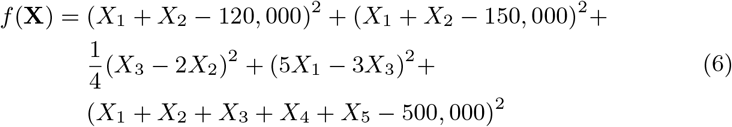

A diverse array of algorithms exist to minimize objective functions for numerical optimization. The relative performance of different algorithms depends on the specific characteristics of the system of equations. Even within the application domain of neuronal census, the choice of the most appropriate algorithm will vary depending on the list of constraints and weights, available computational power, and precision requirements. A comprehensive review of available algorithms is beyond the scope of the present article, and we only mention selected examples instead. For nonlinear constraint optimization, the Levenberg-Marquardt algorithm (Moré 1978), commonly referred to as the “nonlinear least squares method”, may be used to optimize over-determined systems. The Interior point algorithm (Byrd 1999) finds the minimum of a constrained multivariable function and is especially useful for solving large nonlinear programming problems. Simulated Annealing is a popular method for the implementation of functions that search for a global minimum of very complex non-linear objective functions with very large number of optima (Xiang et al. 2013). Among constraint optimization methods suitable for solving linear systems (Coleman and Li 1996; Gill et al. 1981), the Trust-Region-Reflective Algorithm solves the least squares problem with or without equality constraints using Newton’s method. Alternatively, the Interior-point-convex Quadprog Algorithm aims to minimize a quadratic function subject to linear constraints and bound constraints.

In addition to the choice of algorithm, boundary constraints and hyperparameters (e.g. temperature for Simulated Annealing) also need to be set. Since there is often no obvious choice for hyperparameters, exploration of a range of different values may be necessary. Several mathematical packages offer the flexibility of using inbuilt functions with different settings for the algorithms. Optimization of the above example objective function using the interior point algorithm (Byrd 1999) implemented in MATLAB using the fmincon function (Byrd 1999) yields the following results: *X*_1_ = 73, 636; *X*_2_ = 61, 364; *X*_3_ = 122, 727; *X*_4_ = [0, 242, 273]; *X*_5_ = 242, 273 − *X*_4_. A random-seed example of the under-constrained parameter value is *X*_4_ = 121, 161 and *X*_5_ = 121, 112.

## 3 Results

As a pilot study, we applied a variant of the above-described methodology to estimate the population size of each known neuron type in the hippocampal subregion of the dentate gyrus (DG). This required (i) a neuronal classification scheme for the dentate gyrus; (ii) extraction of neuron type count data from the literature; (iii) formatting of literature-extracted data as equations; (iv) optimization.

### 3.1 Classification Scheme

We defined dentate gyrus cell types based on the knowledge base Hippocampome.org, an online repository containing morphological, molecular, and physiological information on neurons of the rodent hippocampal formation (Wheeler et al. 2015). Hippocampome.org classifies neurons primarily by neurotransmitter released and the presence of axons and dendrites in the distinct sub-regions and layers of the rodent hippocampus (Figure 1), and also includes molecular biomarker (Hamilton et al. 2017) and electrophysiological properties (e.g. Lubke et al. 1998) for each type. Hippocampome.org identifies 18 distinct neuron types in DG: 13 with cell bodies exclusively present in a single layer and 5 with cell bodies distributed across two layers. Thus the target unknowns for this neuronal census consist of the population counts for 23 layer-wise types, which we represented here with parameters *X*_1_, *X*_2_, … *X*_23_ (Table 1).

**Fig. 1.**
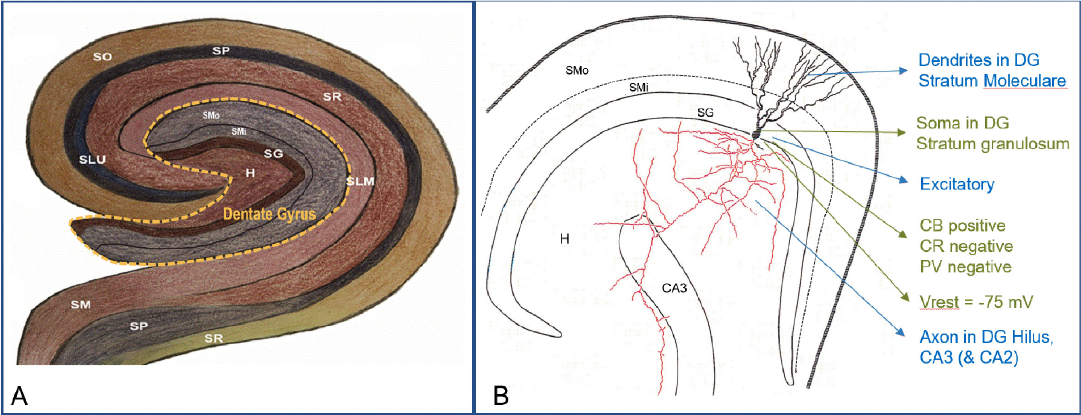
Hippocampome.org neuron type classification. A. Layer organization of the rodent hippocampus, highlighting the dentate gyrus and surrounding regions. B. A dentate gyrus granule cell (cell body and dendrites: black, axon: red) with color-coded properties (neurotransmitter and axonal-dendritic distributions: blue; molecular expression and electrophysiology: green). Label abbreviations: CB: calbindin; CR, calretinin; H, Hilus; PV, parvalbumin; SG, stratum granulosum; SLM, stratum lacunosum-moleculare; SLU, stratum lucidum; SM/SMi/SMo, stratum moleculare (inner/outer); SP, stratum pyramidale; SR, stratum radiatum; Vrest, resting voltage potential.

**Table 1.**
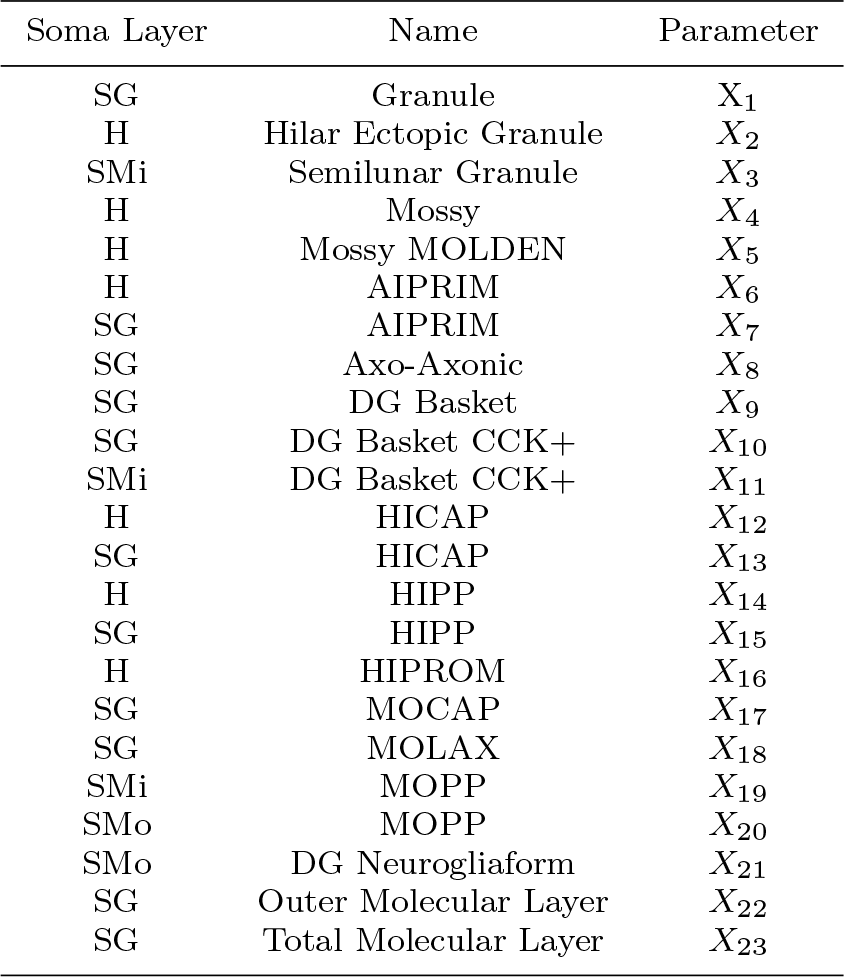
Layer-specific neuron types (left, middle) and parameters representing corresponding counts (right). See legend of Fig. 1 for layer abbreviation definitions.

| Soma Layer | Name | Parameter |
| --- | --- | --- |
| SG | Granule | $X_1$ |
| H | Hilar Ectopic Granule | $X_2$ |
| SMi | Semilunar Granule | $X_3$ |
| H | Mossy | $X_4$ |
| H | Mossy MOLDEN | $X_5$ |
| H | AIPRIM | $X_6$ |
| SG | AIPRIM | $X_7$ |
| SG | Axo-Axonic | $X_8$ |
| SG | DG Basket | $X_9$ |
| SG | DG Basket CCK+ | $X_{10}$ |
| SMi | DG Basket CCK+ | $X_{11}$ |
| H | HICAP | $X_{12}$ |
| SG | HICAP | $X_{13}$ |
| H | HIPP | $X_{14}$ |
| SG | HIPP | $X_{15}$ |
| H | HIPROM | $X_{16}$ |
| SG | MOCAP | $X_{17}$ |
| SG | MOLAX | $X_{18}$ |
| SMi | MOPP | $X_{19}$ |
| SMo | MOPP | $X_{20}$ |
| SMo | DG Neurogliaform | $X_{21}$ |
| SG | Outer Molecular Layer | $X_{22}$ |
| SG | Total Molecular Layer | $X_{23}$ |

### 3.2 Literature Mining

We mined several hundred publications for two kinds of data: (i) ratios of neuron types from studies that morphologically characterized small (i.e. less than 100 cells) populations of neurons; (ii) stereological count estimates of stained neuronal populations. The morphological studies were typically in vitro recording experiments in which neurons were selected non-randomly (i.e. according to explicit target criteria). We assumed the sampling within the bounds of the selection criteria, however, to be uniform. On this assumption, subpopulation ratios within the recorded-from population could be reasonably extrapolated to the region as a whole.

### 3.3 Equation Generation

The sets of neurons as identified by the authors in the mined literature did not directly align with the Hippocampome.org classification scheme. Literature-defined neuron types (literature types) therefore needed to be mapped to Hippocampome.org types. Since each Hippocampome.org type has a unique set of morphological, electrophysiological, and biochemical properties, we translated the description of each literature type into a similarly formalized set of properties, which we then used to match one or more Hippocampome.org types. The literature type could next be assigned the parameter(s) *X*_*i*_ associated with the matching type(s). When a literature type had properties matching multiple Hippocampome.org types, the sum of the parameters representing the corresponding Hippocampome.org types was used.

Here is an example of an equation generated from a morphological classification experiment. One of the mined articles (Ceranik et al. 1997) states: “Neurons from dentate gyrus outer molecular layer were recorded and filled with biocytin for videomicroscopy. 40 neurons were adequately stained. Out of these, 6 neurons were identified as displaced granule cells, 14 neurons had a local axonal arborization that was confined mainly to the OML, 3 projected to the stratum lacunosum moleculare of the CA1 region, and 17 neurons projected to the subiculum via the hippocampal fissure.” In this description, the author defines 4 different groups of neurons having somata in the DG outer molecular layer (SMo in Fig. 1 and Table 1 above). Based on the descriptions of their axons, the groups of 14 and 17 neurons were matched to unique Hippocampome types MOPP and DG Neurogliaform, respectively. This allowed us to construct the equation 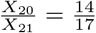, where *X*_20_ and *X*_21_ are the parameters representing the respective outer molecular layer counts of MOPP and Neurogliaform cells. The “displaced granule cells” and “3 project[ing] to the stratum lacunosum moleculare” represented groups with no corresponding Hippocampome types, so similar equations could not be constructed for these groups.

We generated a set of 24 equations (Table 2) using this method. All but one of the equations was extracted from either a stereological calculation or an experiment that morphologically or electrophysiologically classified neurons selected for slice recordings. One equation was based on visual inspection of images of stained slices. While there were more equations than parameters, some of the equations were redundant (multiple studies independently estimated the same quantity). Therefore, the system was both underdetermined and inconsistent.

**Table 2:**
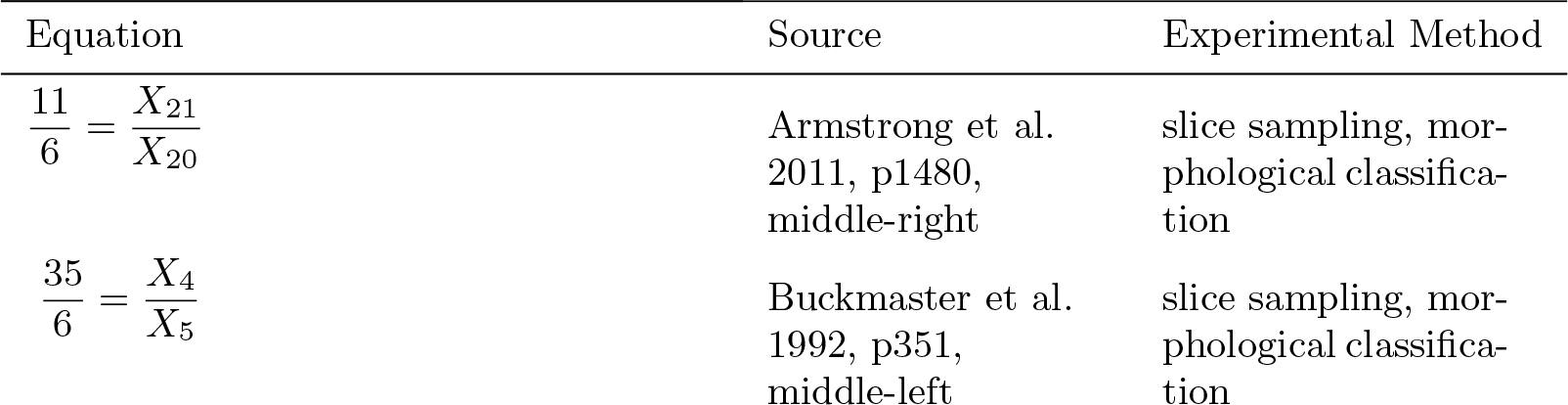
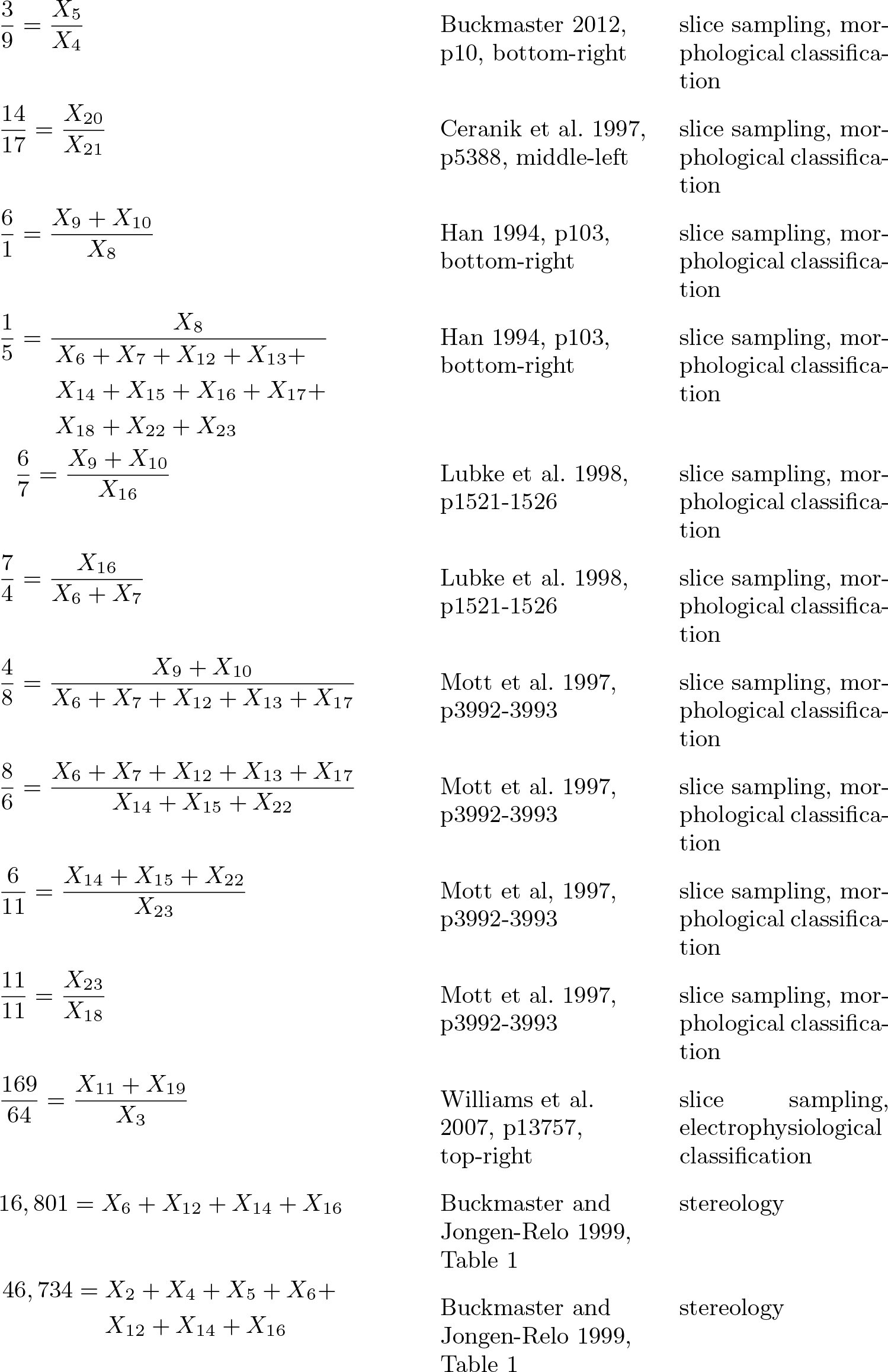
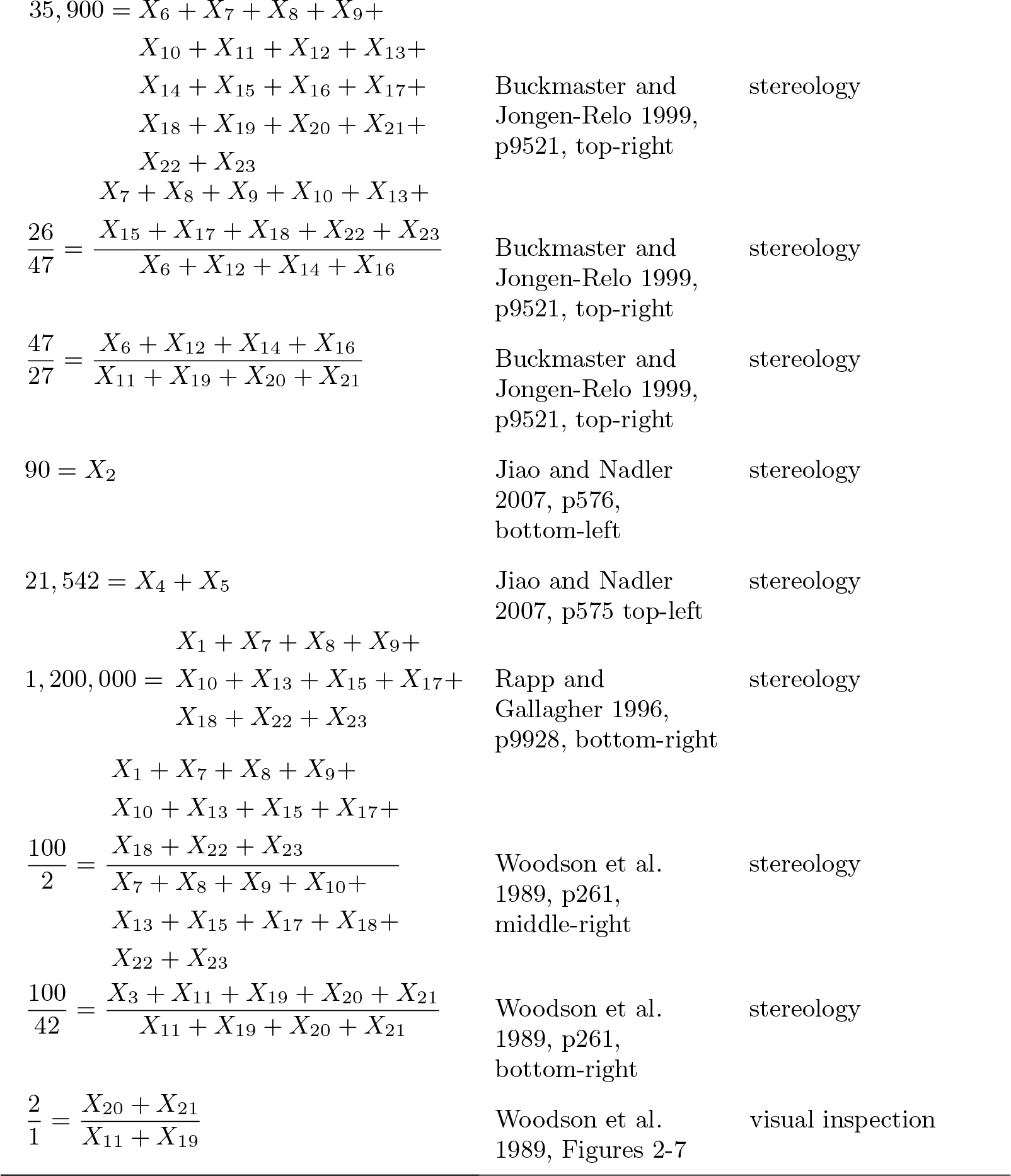
System of equations derived for DG neuron types.

### 3.4 Optimization

An objective function was constructed by summing the weighted, normalized errors associated with each equation, i.e. the absolute value of the difference between left and right sides. Because we considered stereological estimates to be more reliable than morphological type ratios, errors associated with stereological estimates were weighted 5 times those associated with type ratios. Each error term was also normalized by dividing by the sum of parameters having non-zero coefficients in the corresponding equation. This prevented the optimization algorithm from being dominated by errors in equations constraining relatively large populations.

We used simulated annealing (R package GENSA; Xiang et al. 2013) to optimize this objective. We averaged results from 80 runs of simulated annealing, using a temperature setting ranging from 130 to 169.5 in 0.5 unit increments. The results are presented in Table 3. A 95 percent confidence interval is presented for each parameter under the assumption that the results are normally distributed. The coefficient of variation shown is the ratio of the standard deviation of the sample mean to the mean. A lower coefficient of variation corresponds to a parameter with relatively low trial-to-trial variability and thus a hypothetically more accurate estimate.

**Table 3.** Optimization results

| Layer | Name | 95% CI | Mean | Coeff Variation ($\frac{\sigma}{\mu}$) |
| --- | --- | --- | --- | --- |
| SG | Granule | 1,161,163 - 1,175,202 | 1,168,182 | 0.0031 |
| H | Hilar Ectopic Granule | 89 - 90 | 89 | 0.0029 |
| SMi | Semilunar Granule | 1,541 - 1,686 | 1613 | 0.0229 |
| H | Mossy | 17149 - 17,549 | 17,349 | 0.0059 |
| H | Mossy MOLDEN | 4,084 - 4,471 | 4,277 | 0.0231 |
| H | AIPRIM | 3,817 - 4,489 | 4,153 | 0.0413 |
| SG | AIPRIM | 375 - 669 | 522 | 0.1437 |
| SG | DG Axo-axonic | 784 - 906 | 845 | 0.0368 |
| SG | DG Basket | 1,632 - 2,488 | 2,060 | 0.1060 |
| SG | DG Basket CCK+ | 2,282 - 3,083 | 2,682 | 0.0762 |
| SMi | DG Basket CCK+ | 1,761 - 2,365 | 2063 | 0.0747 |
| H | HICAP | 3,210 - 4,256 | 3,733 | 0.0715 |
| SG | HICAP | 105 - 262 | 183 | 0.2183 |
| H | HIPP | 3,194 - 4,292 | 3,743 | 0.0748 |
| SG | HIPP | 144 - 387 | 266 | 0.2335 |
| H | HIPROM | 8,235 - 9,157 | 8,696 | 0.0270 |
| SG | MOCAP | 93 - 266 | 180 | 0.2459 |
| SG | MOLAX | 2,025 - 2,354 | 2,189 | 0.0383 |
| SMi | MOPP | 1,588 - 2,238 | 1,913 | 0.0867 |
| SMo | MOPP | 2,990 - 3,173 | 3,081 | 0.0151 |
| SMo | DG Neurogliaform | 4,515 - 4,763 | 4,639 | 0.0136 |
| SG | Outer Molecular Layer | 88 - 186 | 137 | 0.1825 |
| SG | Total Molecular Layer | 2,257 - 2,631 | 2,444 | 0.0390 |

These results come with several caveats. First, many of the equations come from experiments with an unknown degree of bias. While modern stereological estimates strive to be unbiased, selections of neurons for electrical recording and subsequent staining (the source of our type ratios) may be biased by many factors such as neuronal size, cell longevity in vitro, etc. Second, our algorithm choice (simulated annealing) and hyperparameter choices (temperature and relative weights of error terms) were somewhat arbitrary. In particular, all error terms associated with type ratios were given the same weight. This approach does not capture the differences in reliability among ratios. A weighting scheme based on the sample sizes associated with each ratio could possibly yield more accurate results. Third, due to the constraints on the total numbers of neurons in each layer, optimization will assign a higher average count to each type given a smaller number of types. The presented estimates are therefore likely slightly inflated due to the exclusion of as-yet undiscovered types as well as of types which are too vaguely described for inclusion in Hippocampome.org. Future analyses can account for these neurons by including additional parameters representing unknown types in each layer.

## 4 Conclusions

This work demonstrates that the proposed pipeline of annotation, conversion into equations, and optimization can yield a possible solution to the neuronal census problem. The DG neuron type counts reported here should only be considered preliminary results and were presented for the sole purpose of providing an in-depth illustration of this methodology. Further research must be conducted before finalizing the appropriate choice of algorithms. Exploring multiple algorithms with different combinations of parameter settings is necessary to test and refine the results. It is essential to select the algorithm, hyperparameter settings, and implementation details that yield the most reliable results.

Ultimately, however, the robustness of results will depend on the availability of data and the quality of constraints. Producing more complete and useful results will thus require feeding further constraints to the optimization algorithm. Possible sources of additional constraints include forthcoming results of ongoing experiments, more thorough data mining of existing literature, and assumptions based on domain expert knowledge. Several researchers have reported estimates of different types of cell populations in the brain using stereology, optical fractionator, and other newer methods (Grady et al. 2003; West et al. 1991; Herculano-Houzel et al. 2013; Bezaire et al. 2016; Murakami et al. 2018; Ero et al. 2018). These estimates have been increasing over the years and this growth is widely expected to continue in the foreseeable future.

As a positive side-effect, even an underdetermined system can be useful by identifying the most under-constrained target unknowns in need of additional experimental evidence. Researchers can leverage this information to design specific experiments for revealing the missing information. In summary, constraint optimization is a powerful method that can generate quantitative estimates for the unknown counts of cell types.

## Acknowledgements

This project is supported by grants R01NS39600 and U01MH114829.

## References

1. Armañanzas, R., Ascoli, G. A. (2015). Towards the automatic classification of neurons. Trends in Neuroscience, 38(5), 307–18.

2. Armstrong, C., Szabadics, J., Tamas, G., Soltesz, I. (2011). Neurogliaform cells in the molecular layer of the dentate gyrus as feedforward gammaaminobutyric acidergic modulators of entorhinal-hippocampal interplay. The Journal of Comparative Neurology, 519(8), 1476–1491.

3. Ascoli, G. A., Wheeler, D.W. (2016). In search of a periodic table of the neurons: Axonal-dendritic circuitry as the organizing principle: Patterns of axons and dendrites within distinct anatomical parcels provide the blueprint for circuit-based neuronal classification. BioEssays, 38(10), 969–976.

4. Bezaire, M. J., Raikov, I., Burk, K., Vyas, D., Soltesz, I. (2016) Interneuronal mechanisms of hippocampal theta oscillations in a full-scale model of the rodent CA1 circuit. ELife, https://doi.org/10.7554/eLife.18566.001

5. Bota, M. Swanson, L. W. (2007). The neuron classification problem. Brain Res Rev, 56(1), 79–88.

6. Buckmaster, P. S., Strowbridge, B. W., Kunkel, D. D., Schmiege, D. L., Schwartzkroin, P. A. (1992). Mossy cell axonal projections to the dentate gyrus molecular layer in the rat hippocampal slice. Hippocampus, 2(4), 349–362.

7. Buckmaster, P. S., Jongen-Relo, A. L. (1999). Highly specific neuron loss preserves lateral inhibitory circuits in the dentate gyrus of kainate-induced epileptic rats. Journal of Neuroscience, 19(21), 9519–9529.

8. Buckmaster, P. S. (2012). Mossy cell dendritic structure quantified and compared with other hippocampal neurons labeled in rats in vivo. Epilepsia, 53(Suppl.1), 9–17.

9. Byrd, R.H., Hribar, M.E., Nocedal, J. (1999). An interior point algorithm for large - scale nonlinear programming. SIAM Journal on Optimization, 9(4), 877–900.

10. Ceranik, K., Bender, R., Geiger, J. R., Monyer, H., Jonas, P., Frotscher, M., et al. (1997). A novel type of GABAergic interneuron connecting the input and the output regions of the hippocampus. Journal of Neuroscience, 17(14), 5380–5394.

11. Coleman, T. F. Li, Y. A. (1996). Reflective Newton Method for Minimizing a Quadratic Function Subject to Bounds on Some of the Variables. SIAM Journal on Optimization, 6(4), 10401058.

12. Ecker, J. R., Geschwind, D. H., Kriegstein, A. R., Ngai, J., Osten, P., Polioudakis, D., et al. (2017). The BRAIN initiative cell census consortium: lessons learned toward generating a comprehensive brain cell atlas. Neuron, 96, 542–557.

13. Ero, C.,Gewaltig, M.,Keller, D., Markram, H. (2018). A Cell Atlas for the Mouse Brain. Frontiers in Neuroinformatics, https://doi.org/10.3389/fninf.2018.00084

14. Gill, P. E., Murray, W., Wright, M. H. (1981). Practical Optimization. Academic Press.

15. Grady, M. S., Charleston, J. S., Maris, D., Witgen, B. M., Lifshitz, J. (2003). Neuronal and Glial Cell Number in the Hippocampus after Experimental Traumatic Brain Injury: Analysis by Stereological Estimation. Journal of Neurotrauma, 20(10), 929–941.

16. Hamilton, D. J., White, C. M., Rees, C.L., Wheeler, D. W., Ascoli, G. A. (2017). Molecular fingerprinting of principal neurons in the rodent hippocampus: A neuroinformatics approach. Journal of Pharmaceutical and Biomedical Analysis, 144(10), 269–278.

17. Han, Z. S. (1994). Electrophysiological and morphological differentiation of chandelier and basket cells in the rat hippocampal formation: a study combining intracellular recording and intracellular staining with biocytin. Neuroscience Research, 19(1), 101–110.

18. Herculano-Houzel, S. (2009) The human brain in numbers: a linearly scaled-up primate brain. Frontiers in Human Neuroscience, https://doi.org/10.3389/neuro.09.031.2009.

19. Herculano-Houzel, S., Ribeiro P., Campos L., Valotta da Silva A., Torres L.B., Catania K.C., et al. (2011). Updated neuronal scaling rules for the brains of Glires (rodents/lagomorphs). Brain, Behavior and Evolution, 78, 302–314.

20. Herculano-Houzel, S., Watson, C., Paxinos, G. (2013) Distribution of neurons in functional areas of the mouse cerebral cortex reveals quantitatively different cortical zones. Frontiers in Neuroanatomy, https://doi.org/10.3389/fnana.2013.00035

21. Januszewski, M., Kornfeld, J., Li, P. H., Pope, A., Blakely, T., Lindsey, L. et al. (2018). High-precision automated reconstruction of neurons with flood-filling networks. Nature Methods, 15(8), 605–610.

22. Jiao, Y., Nadler, J. V. (2007). Stereological analysis of GluR2-immunoreactive hilar neurons in the pilocarpine model of temporal lobe epilepsy: correlation of cell loss with mossy fiber sprouting. Experimental Neurology, 205(2), 569–582.

23. Lubke, J., Frotscher, M., Spruston, N. (1998). Specialized electrophysiological properties of anatomically identified neurons in the hilar region of the rat fascia dentata. Journal of Neurophysiology, 79(3), 1518–1534.

24. Morée, J.J. (1978). The Levenberg-Marquardt algorithm: Implementation and theory. Numerical Analys, 630.

25. Mott, D. D., Turner, D. A., Okazaki, M. M., Lewis, D. V. (1997). Interneurons of the dentate-hilus border of the rat dentate gyrus: morphological and electrophysiological heterogeneity. Journal of Neuroscience, 17(11) 3990–4005.

26. Murakami, T. C., Mano, T., Saikawa, S., Horiguchi, S. A., Shigeta, D., et al. (2018). A three-dimensional single-cell-resolution whole-brain atlas using CUBIC-X expansion microscopy and tissue clearing. Nature Neuroscience, 21, 625–637.

27. Plycz, M., Roskams, J., Hill, S., Spruston, N., Meijering, E., et al. (2015). BigNeuron: Large-scale 3D Neuron Reconstruction from Optical Microscopy Images. Neuron, 87(2), 252256.

28. Rapp, P. R., Gallagher, M. (1996). Preserved neuron number in the hippocampus of aged rats with spatial learning deficits. Proceedings of the National Academy of Sciences of the United States of America, 93(18), 9926–9930.

29. Tasic, B., Yao, Z., Smith, K.A., Graybuck, L., Nguyen, T., Bertagolli, D., et al. (2018). Shared and distinct transcriptomic cell types across neocortical areas. Nature, 563(7729), 7278.

30. West, M. J., Slomianka, L., Gundersen, H. J. (1991). Unbiased stereological estimation of the total number of neurons in the subdivisions of the rat hippocampus using the optical fractionator. The Anatomical Record, 231(4), 482–497.

31. Wheeler, D. W., et al. (2015). Hippocampome.org: a knowledge base of neuron types in the rodent hippocampus. Elife, 4.

32. Williams, P. A., Larimer, P., Gao, Y., Strowbridge, B. W. (2007). Semilunar granule cells: glutamatergic neurons in the rat dentate gyrus with axon collaterals in the inner molecular layer. Journal of Neuroscience, 27(50),13756–13761.

33. Woodson, W., Nitecka, L., Ben-Ari, Y. (1989). Organization of the GABAergic system in the rat hippocampal formation: a quantitative immunocytochemical study. The Journal of comparative neurology, 280(2), 254–271.

34. Xiang, Y., Gubian, S., Suomela, B., Hoeng, J. (2013). Generalized Simulated Annealing for Global Optimization: The GenSA Package. The R Journal, 5.

